# Healing of Primary and Recurrent Mouse Ocular Herpetic Disease Following Topical Ocular Treatment with an Engineered Fibroblast Growth Factor-1 (FGF-1) is Associated with Increased Frequency and Function of Corneal Anti-Inflammatory M2 Macrophages

**DOI:** 10.1101/2020.11.25.399360

**Authors:** Nisha R Dhanushkodi, Ruchi Srivastava, Pierre-Gregoire A. Coulon, Swayam Prakash, Soumyabrata Roy, Didier Bagnol, D. David Eveleth, Lbachir BenMohamed

## Abstract

Herpes simplex virus 1 (HSV-1) infects the cornea and caused blinding ocular disease. In the present study, we evaluated whether and how a novel engineered version of fibroblast growth factor-1 (FGF-1), designated as TTHX1114, would reduce the severity of HSV-1-induced primary and recurrent ocular herpes in the mouse model. The efficacy of TTHX1114 against corneal keratopathy was assessed in B6 mice following corneal infection with HSV-1, strain McKrae. Starting day one post infection (PI), mice received TTHX1114 for 14 days. The severity of primary stromal keratitis and blepharitis were monitored up to 28 days PI. Inflammatory cell infiltrating infected corneas were characterized up to day 21 PI. The severity of recurrent herpetic disease was quantified in latently infected B6 mice up to 30 days post-UVB corneal exposure. The effect of TTHX1114 on M1 and M2 macrophage polarization was determined *in vivo* in mice and *in vitro* on primary human monocytes-derived macrophages. Compared to HSV-1 infected non-treated mice, the infected and TTHX1114 treated mice exhibited a significant reduction of primary and recurrent stromal keratitis and blepharitis, without affecting virus corneal replication. The therapeutic effect of TTHX1114 was associated with a significant decrease in the frequency of M1 macrophages infiltrating the cornea, which expressed significantly lower levels of pro-inflammatory cytokines and chemokines. This polarization toward M2 phenotype was confirmed *in vitro* on human primary macrophages. This pre-clinical finding suggests use of this engineered FGF-1 as a novel immunotherapeutic regimen to reduce primary and recurrent HSV-1-induced corneal disease in the clinic.

**IMPORTANCE:** Herpes Simplex Virus type 1 (HSV-1) is a prevalent human pathogen that infects the cornea causing potentially blinding herpetic disease. The present study is the first to demonstrate a reduction of primary and recurrent corneal herpetic disease in mice following engineered FGF-1 (TTHX1114) treatment. This was associated with a decrease of pro-inflammatory M1 macrophages and an increase of anti-inflammatory M2 macrophages infiltrating the corneas of HSV-1 infected mice. The finding in mice were confirmed in humans by showing that in vitro FGF-1 treatment skewed the polarization of primary monocyte-derived macrophages into the anti-inflammatory M2 phenotype. Moreover, FGF-1 treatment appeared to reduce production of anti-inflammatory mediators. This pre-clinical finding support this engineered FGF-1 as a novel immunotherapeutic regimen to reduce primary and recurrent corneal herpetic disease in the clinic.

## INTRODUCTION

With a staggering one billion individuals worldwide currently carrying herpes simplex virus type 1 (HSV-1), herpes remains one of the most prevalent viral infections of the eye (1–7). Ocular herpes infection causes a spectrum of clinical manifestations ranging from blepharitis, conjunctivitis, and dendritic keratitis to disciform stromal edema and blinding stromal keratitis (HSK) (8, 9). In the United States alone, over 450,000 people have a history of recurrent ocular HSV requiring doctor visits, antiviral drug treatments, and in severe cases, corneal transplants (10–12). Despite the availability of many intervention strategies, the global picture for ocular herpes continues to deteriorate (13). Current anti-viral drug therapies (e.g. Acyclovir and derivatives) do not eliminate the virus and reduce recurrent herpetic disease by only ~45% (14). The development of an effective therapy to alleviate ocular disease and heal corneal herpetic scarring would present an unparalleled alternative to anti-viral drugs, as it would be a powerful and cost-effective means to lessen associated blinding ocular herpetic disease (reviewed in (15)).

An intact and fully differentiated corneal epithelium and stroma is critical for proper vision. However, damage and perturbation of the corneal epithelium and stroma is prevalent following exposure to infectious pathogens, such as HSV-1. Ocular infection with HSV-1 can cause eye disease ranging in severity from blepharitis, conjunctivitis, and dendritic keratitis, to disciform stromal edema and necrotizing stromal keratitis. HSV-1 infection of the cornea induces lymphangiogenesis that continues to develop well beyond the resolution of infection. Excessive proteolysis, inflammation and neovascularization, resulting in corneal scarring has been associated with loss of corneal clarity (16). Inflammatory leukocyte-infiltrates the cornea and have been implicated to be essential for corneal neovascularization, an important clinically relevant manifestation of stromal keratitis. An effective medical treatment of vision-threatening corneal herpetic disease is a major unmet clinical challenge.

Multiple pro-angiogenic factors, including the fibroblast growth factor-1, known as FGF-1, are expressed within the cornea following virus clearance. FGF appears to maintain progressive corneal neovascularization following HSV-1 infection; however, treatment with FGF-2 does not appear to increase neovascularization persisting after the peak of disease (17). In the present study, we hypothesized that FGF-1 treatment will: (1) modulate the molecular mechanisms that promote corneal healing and preserved visual acuity in response to primary and recurrent HSV-1 infection; (2) accelerate healing of corneal herpetic disease following primary and recurrent ocular infection with a virulent HSV-1 strain.

Herein, we report that compared to HSV-1 infected non-treated mice, the infected and engineered FGF-1 (TTHX1114) treated mice showed (*i*) an overall resistance to disease and death; (*ii*) a significant decrease in primary stromal keratitis (on days 5, 14, and 21) and blepharitis (on days 7 and 14); and (*iii*) a significant increase in the frequency and function of corneal anti-inflammatory M2 macrophages and a decrease in corneal pro-inflammatory macrophages and inflammatory cytokines. However, eFGF-1 treatment did not affect the number and function of cornea resident T cells nor virus corneal replication. Topical corneal treatment with the eFGF-1, was associated with reduced corneal keratopathy in a mouse model of primary ocular herpes. This pre-clinical finding suggests that inclusion of this engineered FGF-1 as a novel immunotherapeutic regimen may reduce primary and recurrent HSV-1-induced corneal immunopathology in the clinic.

## MATERIALS AND METHODS

### Virus propagation and titration

For virus propagation, rabbit skin (RS) cells (ATCC, Manassas, VA) were grown in Minimum Essential Medium Eagle with Earl’s salts and L-Glutamine (Corning, Manassas, VA) supplemented with 10% fetal bovine serum and 1% penicillin-streptomycin. The HSV-1 laboratory strain McKrae was propagated in RS cells as described previously (20–22) and purified by ultracentrifugation in sucrose gradient and titrated by the plaque assay.

### Mice and infection

Animal protocols were approved by the University of California Irvine’s institutional animal care and use committee (IACUC #19-111). All animals were handled with care according to the guidelines of American Association for Laboratory Animal Science (AALAS). For primary herpes infection, six to eight-week old male and female C57BL6/J mice were purchased from the Jackson Laboratory. The mice were anaesthetized with xylazine (6.6mg/kg) and ketamine (100mg/kg) prior to infection. Both corneas in each mouse was briefly scarified with a 25-gauge needle, tear film blotted, and 1×10^5^ or 5×10^5^ pfu/eye of HSV-1 (strain McKrae) in 2 μL of sterile PBS were inoculated on to the cornea. For herpes reactivation experiments, Wildtype B6 mice were infected with HSV-1 (McKrae 5X10^5^ pfu/eye) after corneal scarification and at day 35 pi, eyes were reactivated by exposure to UV-B radiation for one minutes.

### TTHX1114 treatment

TTHX1114 (N-Met C16S/A66C/C117V FGF1) was prepared as described (Eveleth et al, 2018). During primary herpes infection, HSV-1-infected mice received topical eye treatment with 400 ng/ml TTHX1114 (4ul/eye i.e., 1.6 ng/eye) or equivalent amount of vehicle (PBS) (mock-treatment) from day1 to da 14 days PI (two times/day). During herpes reactivation experiment, one group of mice was mock treated while another group was treated topically with TTHX1114 from day 34 pi for two weeks (two doses each day of 1.6 ng/eye). Unpolarized M0 macrophages were generated from M-CSF-treated primary monocytes and treated with TTHX1114 (0.5 and 3 ng/ml) for 24 hr. M1- and M2-polarized macrophages were then generated by stimulation with IFN-γ and IL-4, respectively (see additional details below).

### Corneal herpetic disease scoring

Mice were monitored for ocular herpes infection and disease progression. To examine corneal inflammation and cloudiness, pictures were taken at several time points with a Nikon D7200 camera. Mice were scored on days 5, 7, 10,14, 21, 28 for pathological symptoms of keratitis and blepharitis after infection with 2X 10^5^ pfu/eye of HSV-1 McKrae, mock/treated with TTHX1114 from day 1 p.i. Stromal keratitis was scored as 0-no disease; 1-cloudiness, some iris detail visible; 2-iris detail obscured; 3-cornea totally opaque; and 4-cornea perforation. Blepharitis was scored as 0-no disease; 1-puffy eyelids; 2-puffy eyelids with some crusting; 3-eye swollen shut with severe crusting; and 4-eye completely swollen shut.

### Quantification of infectious virus

Tears were collected from both eyes using a Dacron swab (type 1; Spectrum Laboratories, Los Angeles, CA) on days 3, 5 and 7 pi. Individual swabs were transferred to a 2mL sterile cryogenic vial containing 1ml culture medium and stored at −80°C until further use. The HSV-1 titers in tear samples were determined by standard plaque assays on RS cells as previously described (18). Eye swabs (tears) were analyzed for viral titers by the plaque assay. RS cells were grown to 70% confluence in 24-well plates. The transfer medium in which eye swabs were stored in was added after appropriate dilution at 250 μl per well in 24-well plates. Infected monolayers were incubated at 37°C for 1 h and were rocked every 15 min for viral adsorption and then overlaid with medium containing carboxymethyl cellulose. After 48 hours of incubation at 37°C, cells were fixed and stained with crystal violet, and viral plaques and counted under a light microscope. Positive controls were run with every assay using previously tittered laboratory stocks of McKrae.

### Monocyte-derived macrophage culture

PBMC were isolated from 20 mL of donor blood by gradient centrifugation using a leukocyte separation medium (Fisher Scientific, Waltham, MA). The cells were washed in PBS and re-suspended in complete culture medium consisting of RPMI-1640 medium containing 10% FBS (Bio-Products, Woodland, CA) supplemented with 1x penicillin/L-glutamine/streptomycin, 1x sodium pyruvate, 1x non-essential amino acids. Monocytes were isolated from PBMC by adherence to culture plate wells for 1 hr. M0 macrophages were cultured and generated from M-CSF(20ng/ml) −treated primary monocytes for 7 days. M1- and M2-polarized macrophages were then generated by stimulation of unpolarized M0 macrophages with IFN-γ(20ng/ml) and IL-4 (20ng/ml) for 48 hours, respectively.

### Immunohistochemistry and confocal microscopy

Corneas were excised after cardiac perfusion with cold PBS of deeply anesthetized mice. The corneas were treated in 4% paraformaldehyde (PFA) for 30 min at 4° C followed by three 15 min wash in PBS in 0.1% Triton X-100 (Sigma) at room temperature (RT). The corneas were blocked with 10% fetal bovine serum overnight at 4° C. For whole corneal staining, the corneas were incubated overnight with a cocktail of rabbit anti-mouse A488 conjugated anti-mouse LYVE-1 (Abcam), PE-conjugated anti-mouse CD31 (Millipore), in PBS with 0.2% Trition X-100. The corneas were washed 5 times, 30 min per wash in 0.1% Trition X-100 PBS at RT, and mounted on a glass slide after making radial cuts. Images were captured on the BZ-X710 All-in-One fluorescence microscope (KEYENCE Corporation of America, Itasca, IL).

### Flow cytometry

C57BL/6 mice infected with HSV-1 (McKrae 2X10^5^ PFU /eye) were treated with TTHX1114 (1.6ng /eye twice daily) and at day 5, 7, 10, 14 and 21 p.i. corneas were pooled (n=6 per group) and stained for FACS analysis. Mice were euthanized at various times p.i. and harvested corneas were digested with collagenase III (5mg/ml) in RPMI 1640 containing 10% fetal bovine serum (FBS), 1% antibiotic/antimycotic, and gentamicin at 37° C. Cornea were dissociated with a 3-mL syringe-plunger head in the presence of media. Cell suspensions were passed through a 40-micron filter before staining. Single cell suspensions were labeled with the following fluorochrome-conjugated monoclonal antibodies: anti-mouse CD45, CD3, CD4, CD8, CD69 GzmB, CD11b, CD11c, F4/80, CD206. For surface staining, mAbs were added against various cell markers to a total of 1 x10^6^ cells in phosphate-buffered saline (PBS) containing 1% FBS and 0.1% sodium azide (fluorescence-activated cell sorter [FACS] buffer) and left for 45 min at 4°C. For intracellular/intranuclear staining, cells were first treated with cytofix/cytoperm (BD Biosciences) for 30 min. Upon washing with Perm/Wash buffer, mAbs were added to the cells and incubated for 45 min on ice in the dark, washed with Perm/TF Wash, FACS buffer and fixed in PBS containing 2% paraformaldehyde. Labeled cells were suspended in 1% BSA in PBS and analyzed using a BD Fortessa flow cytometer.

For flow cytometry staining of human monocytes-derived macrophage (MDM) macrophages were harvested after treatment with accutase for 30 min and vigorous washing with cold PBS. The macrophage suspension was stained with anti-human CD45, CD11b, Cd14, CD68, CD80, CD64, CD163, CD206 antibodies respectively. Labeled cells were suspended in 1% BSA in PBS and analyzed using a BD Fortessa flow cytometer.

### Luminex assay

Corneal lysates or cell supernatants were assayed for cytokines IFN-γ, IL-1α, IL-1β, IL-2, IL-4, IL-5, IL-6, IL-10, IL-12-p40, IL-12-p70, IL-15, IL-17, IP-10, GM-CSF and TNF-α using the Luminex kit according to the manufacturer’s instructions (Milliplex Multiplex Assays with Luminex, Millipore Sigma, Danvers, MA). Samples were assayed using the Luminex assay system (Magpix).

### Statistical analyses

Data for each assay were compared by analysis of variance (ANOVA) and Student’s *t* test using GraphPad Prism version 5 (La Jolla, CA). Differences between the groups were identified by ANOVA and multiple comparison procedures, as we previously described (19). Data are expressed as the mean + SD. Results were considered statistically significant at *p* < 0.05.

## RESULTS

### 1. FGF-1 topical ocular treatment reduced primary ocular herpes stromal keratitis and blepharitis in mice, independent of virus replication

We first investigated the effect of the Engineered Fibroblast Growth Factor-1 (FGF1 also designated as TTHX1114, structure illustrated in **Fig. 1A**) on corneal keratopathy in C57BL/6 (B6) mice infected ocularly with 1 x 10^5^ or 5 x 10^5^ PFUs of HSV-1 (strain McKrae). Starting day one post-infection (PI) B6 mice received daily topical ocular treatment with TTHX1114 twice daily (8 hour. intervals) for 14 days (**Fig. 1B**). The efficacy of TTHX1114 on primary corneal infection and disease was tested at an initial dose of 1.6 ng/eye, twice a day. The severity of primary stromal keratitis and blepharitis was monitored on days 2, 5, 7, 10, 14, 21 and 28 PI (**Fig. 1B**). HSV-1 replication in cornea was also determined at 2, 5, 7, 10 days PI (**Fig. 1B**). As shown in the representative corneal pictures in **Fig. 1C**, compared to HSV-1 infected vehicle-treated mice (*lower panel*), the infected and TTHX1114-treated mice (*top panel*) showed a significant decrease of corneal herpetic disease. The most significant decrease in primary stromal keratitis was recorded on days 14 and 21 (*P* = 0.04 and *P* = 0.02, respectively, **Fig. 1D**). A significant decrease in blepharitis was recorded on days 7 and 14 (*P* = 0.02 and *P* = 0.04, respectively, **Fig. 1E**). However, there was no significant effect observed with TTHX1114 on mouse survival following HSV-1 infection (**Fig. 1F**). No significant effect of TTHX1114 on corneal virus replication was detected (*not shown*). The effect of TTHX1114 treatment was recorded at both high dose 5 x 10^5^ PFUs and low 1 x 10^5^ dose of HSV-1, on blepharitis as early as day 5 post-treatment (**Fig. 2A**) and on keratitis at day 7 post-treatment (**Fig. 2B**). Immunohistochemistry and FACS analysis were carried out to assess if FGF-1 treatment can affect lymphangiogenesis and lymphocyte infiltration in HSV-1-infected mice. The significant reduction in primary corneal keratopathy following TTHX1114 treatment was not associated with a reduction in lymphangiogenesis (**Supplemental Fig. 1**).

**Figure 1.**
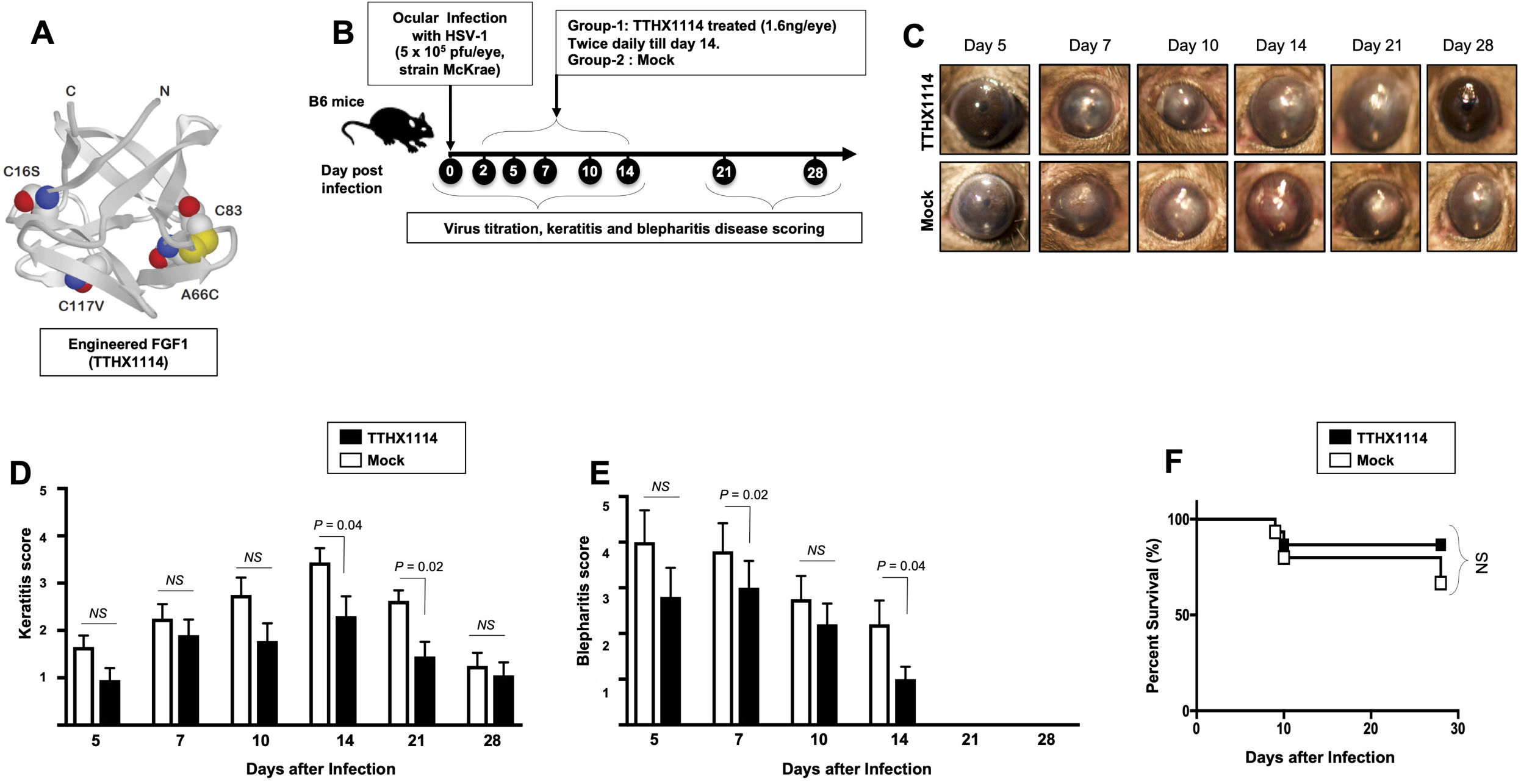
Effect of FGF-1 treatment on corneal disease during acute corneal HSV-1 infection: (**A**) Structure of FGF-1, an engineered FGF-1 known to aid in corneal epithelial wound healing. (**B**) Experimental plan to assess the effect of FGF-1 topical eye treatment in HSV-1 (McKrae) infected B6 mice is shown. Mice were mock treated/ treated with FGF-1 (1.6 ng/eye, twice a day) from day1 post-infection with HSV-1 McKrae (5X10^5^/eye). Mice were scored at day 5, 7, 10,14, 21, 28 for pathological symptoms of Keratitis and blepharitis after infection with 5X 10^5^ pfu/eye of HSV-1 McKrae, mock/treated with FGF-1 from day1 p.i. Stromal keratitis was scored as 0-no disease; 1-cloudiness, some iris detail visible; 2-iris detail obscured; 3-cornea totally opaque; and 4-cornea perforation. Blepharitis was scored as 0-no disease; 1-puffy eyelids; 2-puffy eyelids with some crusting; 3-eye swollen shut with severe crusting; and 4-eye completely swollen shut. Keratitis score and Blepharitis score in mice mock/ treated with FGF-1 during corneal HSV-1 infection. (**C**) Representative eye pictures of mice mock treated/ treated with FGF-1 from day1 after infection with HSV-1 McKrae. Graph showing corresponding keratitis (**D**) and blepharitis score (**E**) at day 7, 10 14 p.i. Data represent the mean score from experiment with a total of 10 mice per group. (**F**) Survival plot of mice mock treated/ treated with FGF-1 (1.6 ng/eye) twice a day from day1 post-infection with HSV-1 McKrae (5X10^5^/eye).

**Figure 2:**
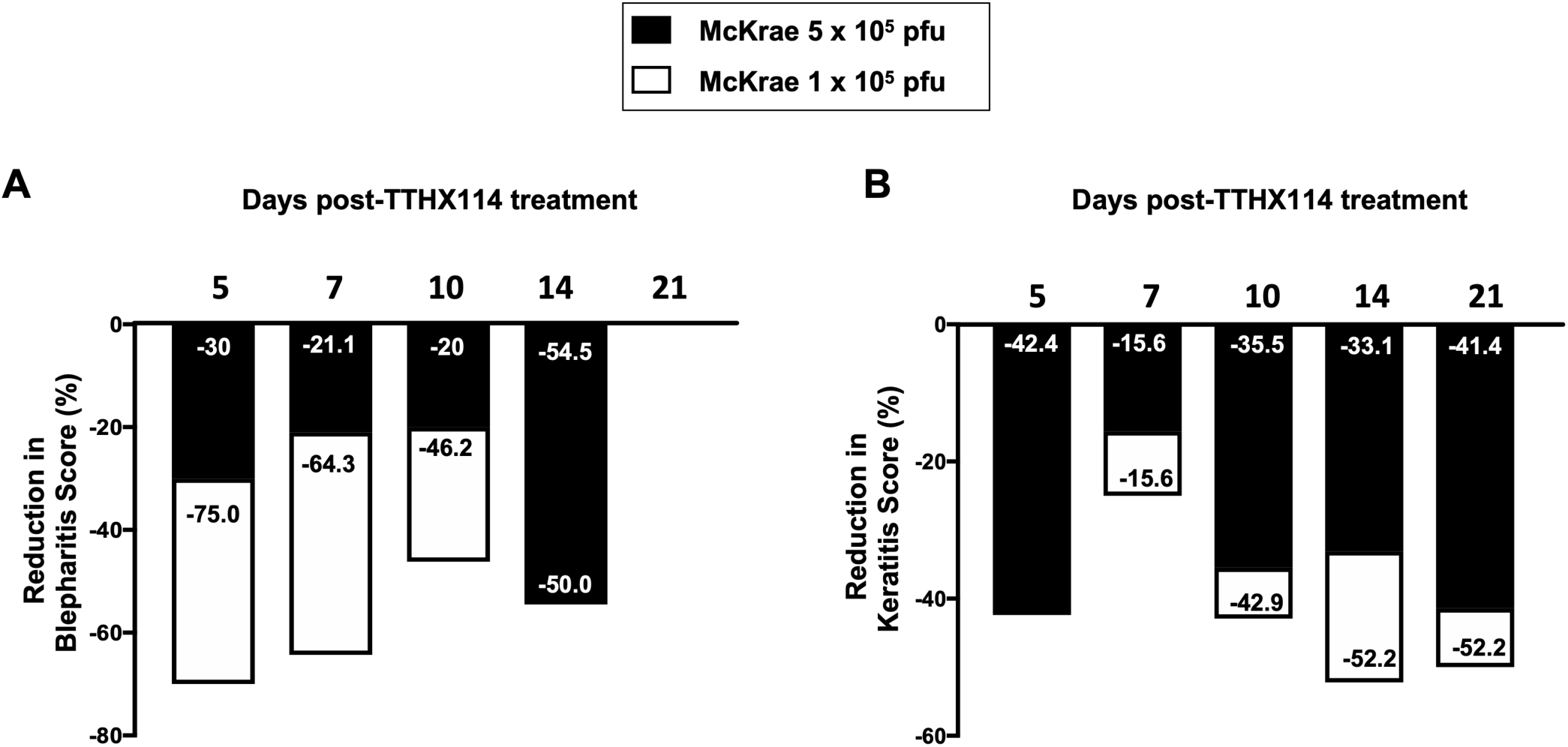
Efficacy of an engineered FGF-1, TTHX1114, on the reduction of severity of primary corneal herpetic disease: Mice were treated, or mock treated, with TTHX1114 (1.6 ng/eye, twice a day) from day1 post-infection with HSV-1 McKrae (1 x 10^5^ and 5 x 10^5^ pfu/eye). Mice were scored for pathological symptoms of blepharitis and Keratitis after infection at day 5, 7, 10, 14, 21 and 28 −days post-infection. Stromal keratitis was scored as 0-no disease; 1-cloudiness, some iris detail visible; 2-iris detail obscured; 3-cornea totally opaque; and 4-cornea perforation. Blepharitis was scored as 0-no disease; 1-puffy eyelids; 2-puffy eyelids with some crusting; 3-eye swollen shut with severe crusting; and 4-eye completely swollen shut. Bar graph illustrating the percentage reduction in blepharitis (**A**) and keratitis (**B**) scores between mice infected with HSV-1 McKrae 1 x 10^5^ (white boxes) or 5 x 10^5^ pfu/eye (black boxes) untreated vs following treatment with TTHX1114. The reduction in blepharitis and keratitis score following treatment with TTHX1114 was calculated as percentage of mock treated group for each scoring day. In A, the efficacy of TTHX1114 in decreasing the blepharitis score at day 14 is identical in mice infected with a low (1 x 10^5^) or high (5 x 10^5^ pfu/eye) HSV-1 McKrae strain titer. The reduction in blepharitis score is otherwise greater at all other time points in mice infected with HSV-1 McKrae 1 x 10^5^ compared to 5 x 10^5^ pfu/eye. In contrast, the reduction in the keratitis score is comparable in mice infected with HSV-1 McKrae 1 x 10^5^ or to 5 x 10^5^ pfu/eye (B).

These results demonstrate that topical cornea treatment with FGF-1 (TTHX1114) is associated with reduced primary corneal keratopathy in the B6 mouse model of primary ocular herpes, independent of viral shedding or lymphangiogenesis.

### 2. FGF-1 topical ocular treatment reduced recurrent herpetic disease in the mouse model of UVB-induced virus reactivation

We next evaluated the effect of TTHX1114 on recurrent herpes infection and disease using the B6 mouse model of UVB induced reactivation (**Fig. 3A**). In this model, the cornea of B6 mice were infected with HSV-1 with scarification and virus reactivation was provoked at day 35 PI in latently infected mice, using a 60 seconds corneal UV-B radiation, immediately followed with topical treatment with TTHX1114 for two weeks. One group of mice was treated topically with TTHX1114 (*n* = 26) from day 34 p.i for two weeks (two doses each day) while another group of mice was mock treated (control *n* = 26). The efficacy of TTHX1114 on recurrent herpetic disease was tested at an initial dose of 1.6 ng/eye, twice a day on the severity of recurrent stromal keratitis monitored daily for 30 post-UVB exposure (**Fig. 3A**). HSV-1 reactivation in the cornea was also determined 10 days PI (**Fig. 3A**). As shown in the representative corneal pictures in **Fig. 3B**, compared to HSV-1 infected non-treated mice (*lower panel*), the infected and TTHX1114-treated mice (*top panel*) showed a significant decrease of recurrent corneal herpetic disease. The most significant decrease in recurrent stromal keratitis was recorded on days 10 and 12 post-UVB-induced reactivation (*P* = 0.04 and *P* = 0.02, respectively, **Figs. 3C–3F**. Further comparison of the distribution of recurrent herpetic disease duration, following UVB induced reactivation, between TTHX1114-treated and mock-treated showed a significantly higher number of eyes displaying recurrent herpetic disease for more than 10 days in the mock-treated compared to TTHX1114-treated group of mice (**Fig. 3G**). However, there was no significant effect of TTHX1114 on virus shedding detected in the cornea (*data not shown*).

**Figure 3.**
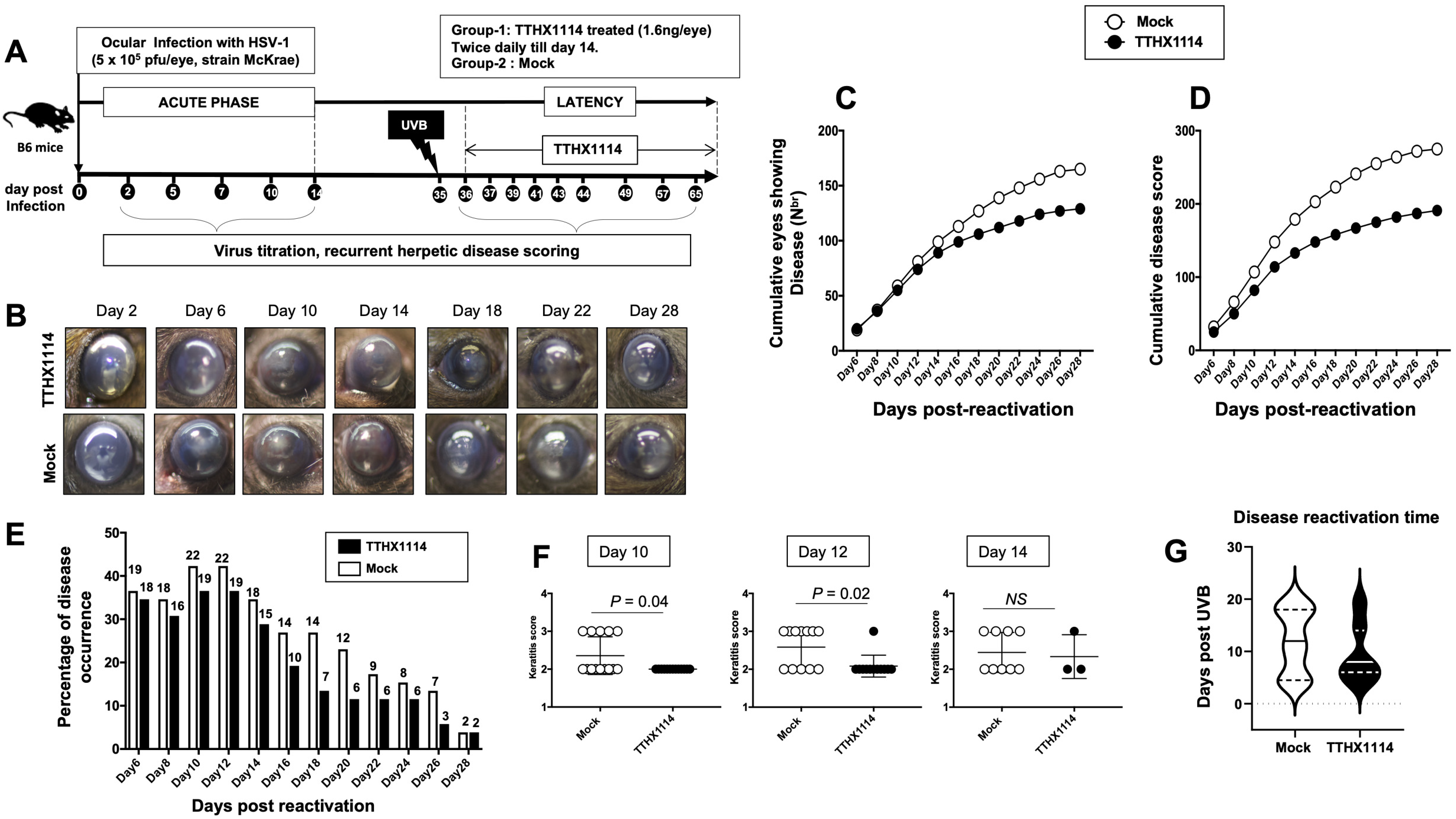
Effect of FGF-1 treatment on recurrent ocular herpes mouse model of UVB-induced herpes reactivation: Wildtype B6 mice were infected with HSV-1(McKrae 5 x 10^6^ pfu/eye and at day 35 p.i, eyes were reactivated by exposure to UV-B radiation for one minute. One group of mice was mock treated (n=26) while another group was treated topically with FGF-1 (n=26) from day 34 p.i for two weeks (two doses each day). (**A**) Experimental plan for testing the effect of FGF-1 on mouse model of ocular herpes reactivation. (**B**) Representative mouse eye picture of herpes UV-B reactivated mice treated with FGF-1. (**C**) Cumulative number of eyes with reactivated disease in mock and FGF-1 treated mice group from day6 to day 28 post-reactivation is shown. (**D**) Graph showing cumulative disease score in mock and FGF-1 treated mice group from day6 to day 28 post-reactivation. (**E**) Percentage of disease occurrence in mock treated and FGF-1 treated mice group from day 2 to day 28 post-reactivation. Number above each bar represents the number of eyes showing disease out of 52 eyes. (**F**) Graph showing keratitis score (more than 2) in mock and FGF-1 treated at day10, day12 and day14 post-reactivation. (**G**) Duration of recurrent corneal herpetic disease in HSV-1 infected mice following treatment with FGF-1. The violin plot illustrates the distribution of disease duration post UV-B radiation in days between mock and TTHX1114 treated groups. Note the higher number of eyes displaying disease for more than 10 days in the mock group compared to TTHX1114 treated mice. Only eyes with disease score above 2 for both mock and treated groups were included in this analysis.

These results demonstrate that topical cornea treatment with the FGF-1 (TTHX1114) is associated with reduced recurrent corneal herpetic disease in the B6 mouse model of UVB induced reactivation independent of the level of virus shedding in the cornea.

### 3. The reduction of corneal herpetic keratopathy following FGF-1 treatment was associated with a decrease of cornea-resident pro-inflammatory M1 macrophages

Since macrophages appeared to be an important inflammatory cell infiltrate in the corneas following ocular herpes infection (20–22), we next assessed the effect of TTHX1114 treatment on the infiltration and function of inflammatory immune cells and determined its correlation with reduction of corneal keratopathy following TTHX1114 treatment in infected corneas. B6 mice were infected with HSV-1 (McKrae 2 x 10^5^ PFU /eye) and then treated with TTHX1114 (1.6 ng /eye twice daily) or left untreated as controls (*mock*). On days 2, 5, 8, 14 and 21, the corneas were harvested, pooled (6 corneas per group) and stained for total CD45^+^CD11b^+^F4/80^+^ macrophages and analyzed by FACS assay using the gating strategy showed in **Supplemental Fig. 2**. As shown in **Figs. 4A, 4B and 4C**, similar frequencies of total CD45^+^CD11b^+^F4/80^+^ macrophages were detected in mouse corneas treated or untreated with TTHX1114. We observed decreased pro-inflammatory Ly6c^high^F4/80^+^CD11b^+^ macrophages on day7, day14 following TTHX1114 treatment (**Figs. 4C**). In addition, we observed a trend towards increased anti-inflammatory CD206^+^F4/80^+^CD11b^+^ macrophages M2 on day7 following TTHX1114 treatment (**Figs. 4C**). These results correlate with a decreased inflammatory cytokine profile in mouse cornea upon TTHX1114 treatment during herpes infection (**Fig. 5**). While a slight increase in CD8^+^ T cells was seen in the cornea of TTHX1114 treated group, no significant effect of TTHX1114 treatment was detected on the frequency and activation of cornea-resident CD4^+^ T cells (**Fig. 6**).

**Figure 4.**
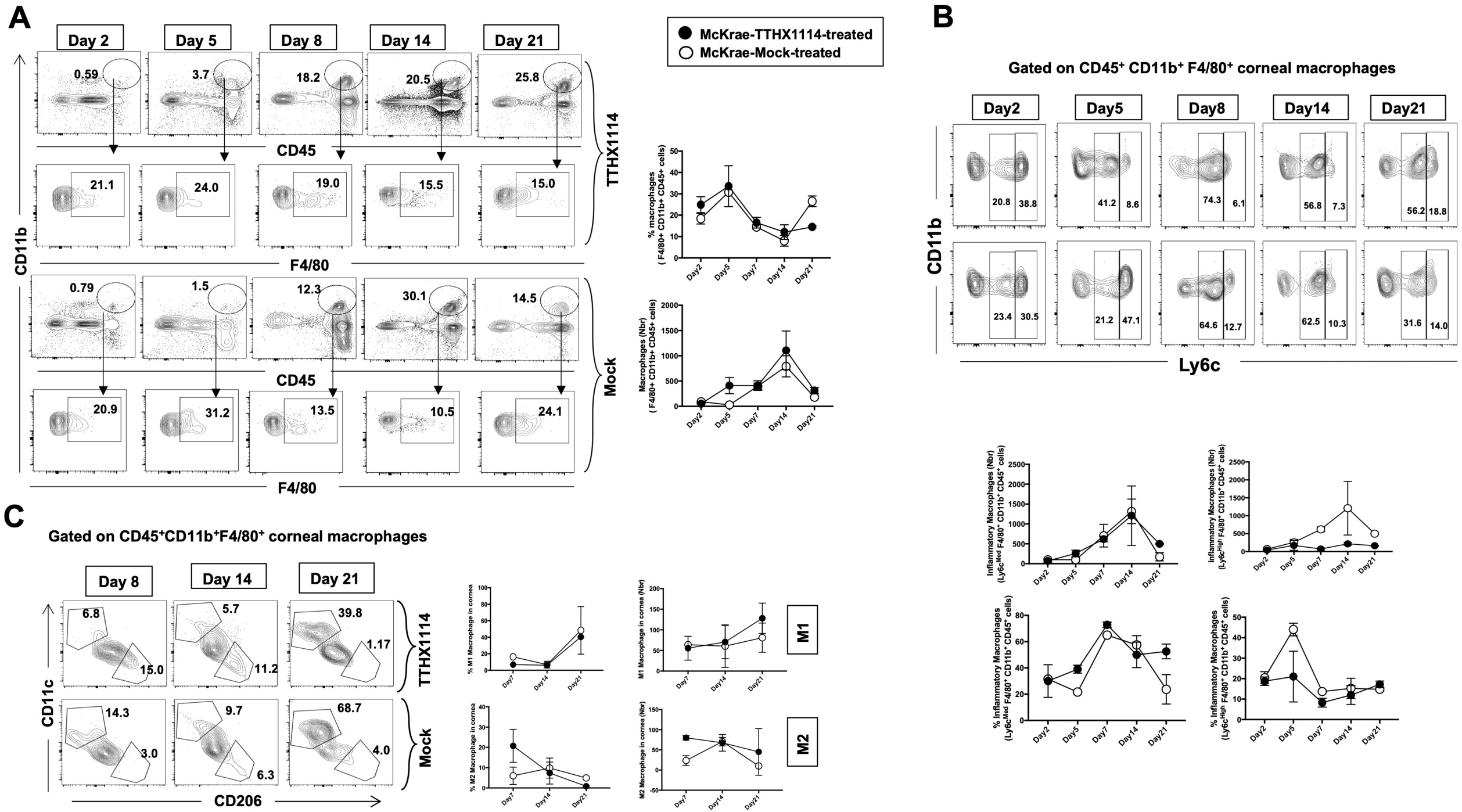
Effect of FGF-1 on the infiltration of M1/M2 macrophages in the cornea of HSV-1 infected B6 mice: FACS analysis was carried out to assess the effect of FGF-1 treatment on inflammatory immune cells infiltration in infected corneas. C57BL/6 mice infected with HSV-1 (McKrae 5X10^5^ PFU /eye) were treated withFGF-1 (1.6ng /eye twice daily) and at day 2, 5, 7, 14 and 21, corneas were pooled (6 per group) and stained for FACS analysis. (**A**) Panel showing FACS plots for macrophages in mouse corneas. Graph (Right panel) showing corresponding average percentage of F4/80^+^CD11b^+^ macrophages in FGF-1 post ocular HSV-1 infection. At day 7, 14 and 21 PI corneas were pooled (6 per group) and stained for FACS analysis. (**B**) FACS plot showing percentage of inflammatory Ly6c^high^F4/80^+^CD11b^+^ and Ly6c^Med^F4/80^+^CD11b^+^ macrophages in FGF-1 post ocular HSV-1 infection. Graph (lower panel) showing corresponding average percentage of Ly6c^high^F4/80^+^CD11b^+^ and M2 Ly6c^Med^F4/80^+^CD11b^+^ inflammatory macrophages in FGF-1 post ocular HSV-1 infection. (**C**) FACS plot showing percentage of M1 (CD11c^+^F4/80^+^CD11b^+^) and M2 (CD206^+^F4/80^+^CD11b^+^) macrophages in FGF-1 post ocular HSV-1 infection. Graph (Right panel) showing corresponding average percentage of M1 (CD11c^+^F4/80^+^CD11b^+^) and M2 (CD206^+^F4/80^+^CD11b^+^) macrophages in FGF-1 post ocular HSV-1 infection.

**Figure 5.**
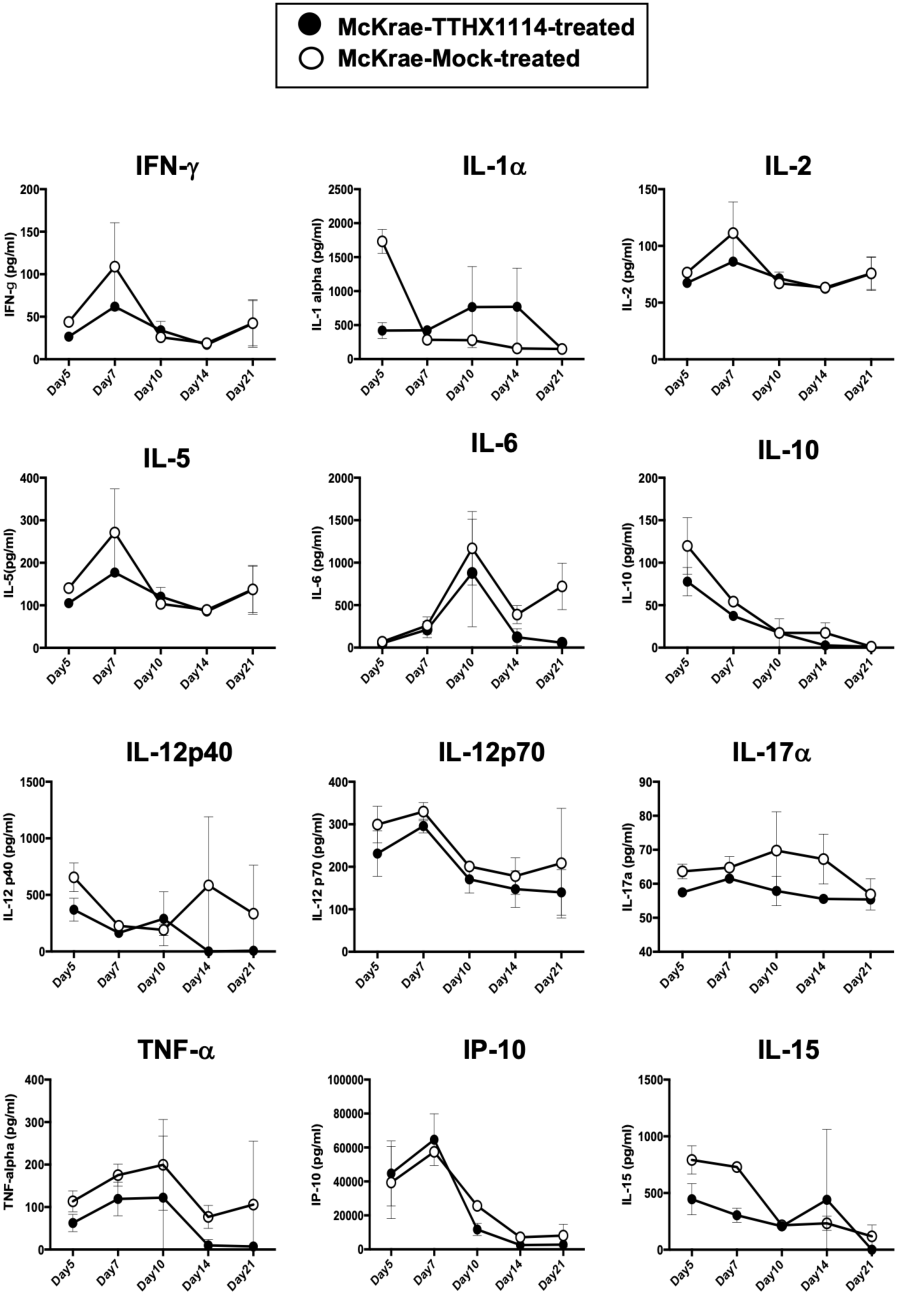
Effect of FGF-1 on the production of inflammatory cytokine in mouse cornea following ocular HSV-1 infection: C57BL/6 mice infected with HSV-1 (McKrae 5X10^5^ PFU /eye) were treated with FGF-1 (1.6ng /eye twice daily) and at day 2, 5, 7, 14 and 21, corneas were pooled (6 per group) and lysates were analyzed by luminex for inflammatory cytokine profile. **(A)** Inflammatory cytokines (IFN-γ, IL-1α, IL-2, IL-5, IL-12p40, IL-12p70, IL-15, IL-17α, IP-10, TNF-α,) levels in the corneal lysates of herpes infected mice treated with TTHX1114 (black circle) compared with mock treated (open circle).

**Figure 6.**
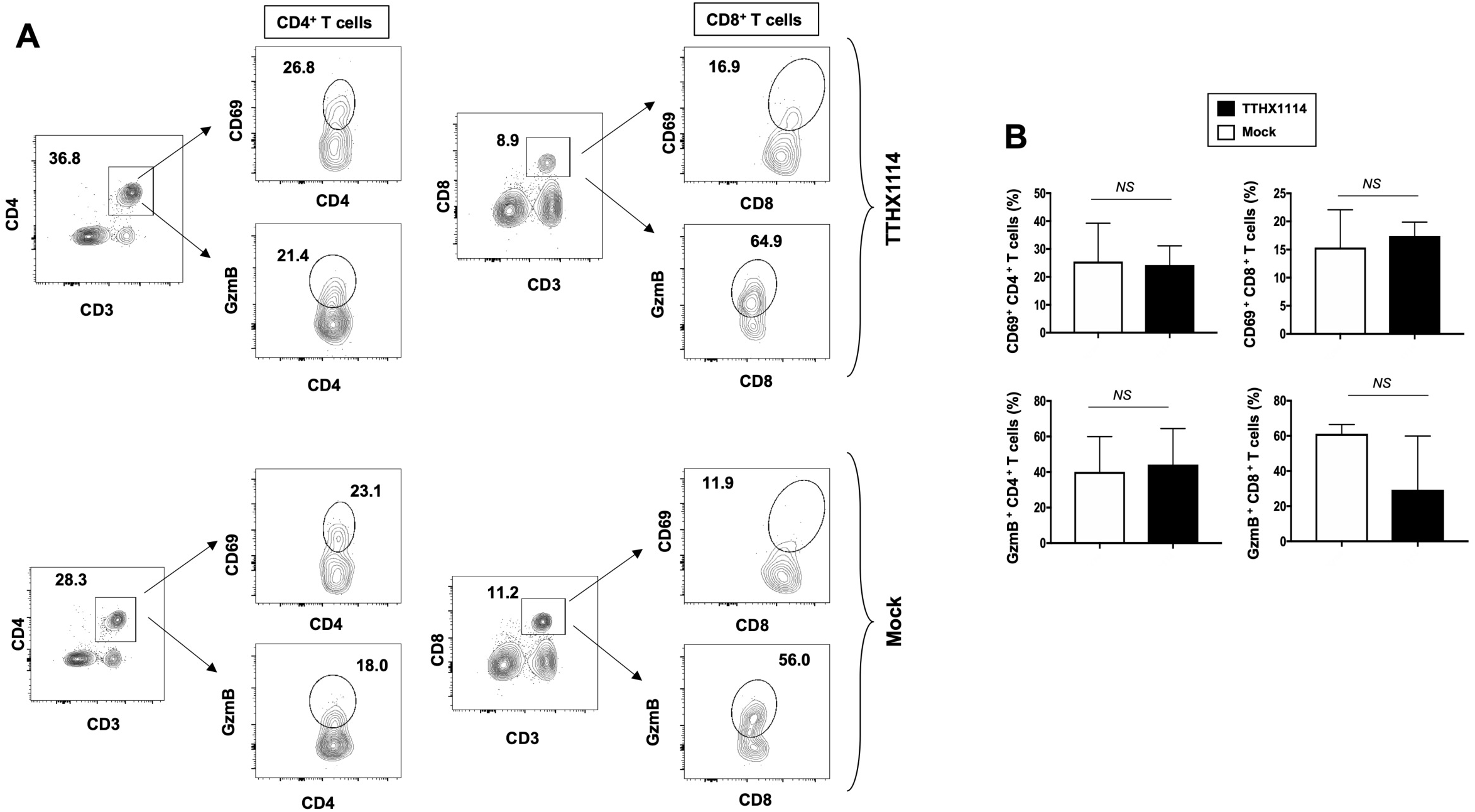
Effect of FGF-1 on lymphocyte activation in mouse cornea during HSV-1 infection: FACS analysis was carried out to assess the effect of FGF-1 treatment on CD4, CD8 T cells activation. C57BL/6 mice infected with HSV-1 (McKrae 5X10^5^ PFU /eye) were treated with FGF-1 (1.6ng /eye twice daily) and at day 8, corneas were pooled (8 per group) and stained for FACS analysis. (**A**) Panel showing FACS plots for CD69 (activation marker), (cytotoxic granular protein) expression in CD4^+^ T cells and CD8^+^ T cells in mouse corneas. (**B**) Graph showing corresponding average percentage of CD69^+^ CD4^+^, GzmB^+^ CD4^+^, CD69^+^ CD8^+^, GzmB CD8^+^ T cells in cornea of B6 mice treated with FGF-1 post ocular HSV-1 infection. Statistical analysis carried out using student’s t test.

These results indicate that reduction of corneal keratopathy following FGF-1 treatment was associated with an alteration in the ratio of cornea-resident M1/M2 macrophages infiltrating the mouse cornea infected with HSV-1. However, there was no association with infiltration nor stimulation of cornea-resident CD4^+^ and CD8^+^ T cells.

### 4. FGF-1 treatment skews polarization of human monocyte into M2 macrophages that produce anti-inflammatory cytokines/chemokines

Based on the mouse results above demonstrating the effect of TTHX1114 on cornea-resident M1/M2 macrophages, we next determined whether this effect would be confirmed on human M1/M2 macrophages. Human unpolarized M0 macrophages were generated from M-CSF-treated primary blood-derived monocytes and then either left untreated or treated with TTHX1114 at 0.5 and 3 ng/mL respectively for 24 hours. The M1- and M2 macrophages were subsequently generated following stimulation with either IFN-γ or IL-4, respectively (**Figs. 7A** and **7B**, *top panels*). The representative images of M1 and M2 macrophages untreated or treated with TTHX1114 showed a different distribution and texture (**Figs. 7A** and **7B**, *middle panels*). **Supplemental Fig. 3** shows expression of M1 and M2 polarization markers in human monocyte-derived macrophages.

**Figure 7.**
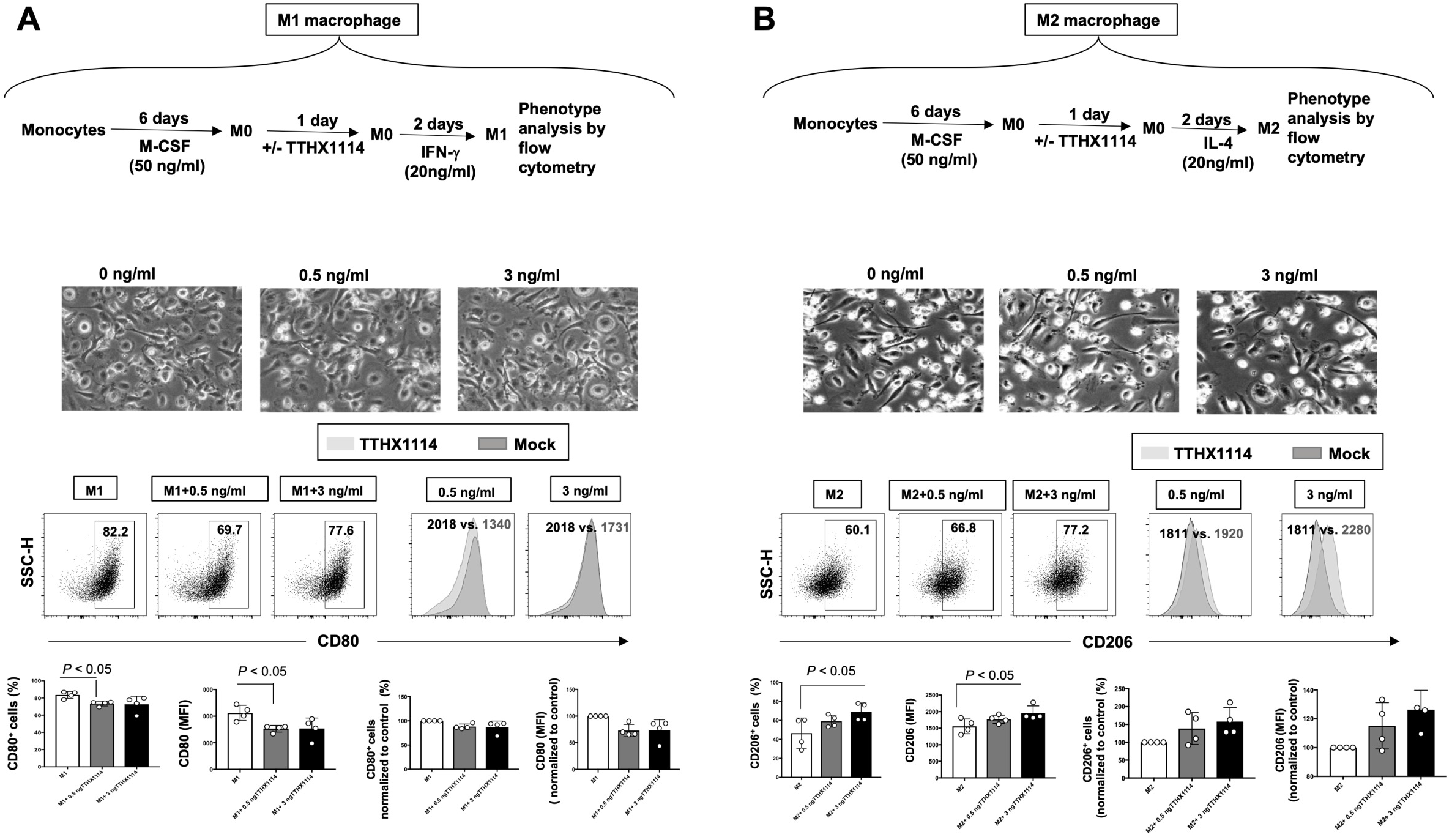
Effect of FGF-1 on M1 and M2 polarization from human monocyte-derived macrophages: Unpolarized M0 macrophages were generated from M-CSF-treated primary monocytes and treated with FGF-1 (0.5 and 3 ng/ml) for 24 hr. M1- and M2-polarized macrophages were then generated by stimulation with IFN-g and IL-4, respectively. (**A**) Experimental plan as shown (top panel). Representative images of M1 and M2 macrophages treated with FGF-1 (0.5 and 3 ng/ml). CD80 and CD64 levels in M1 macrophages treated with FGF-1 were compared by flow cytometry. Dot plots and histograms depict the results obtained in one representative donor; the grey histogram represent mock treated M1 macrophage, and the blue histogram represent the fluorescent profile of FGF-1 treated M1 macrophage stained with the indicated antibodies; the percentage and MFI (mean fluorescence intensity) of positive cells is indicated. The graph (lower panel) represents mean results from at least three different donors. (**B**) Experimental plan as shown (top panel). CD163 and CD206 levels in M2 macrophages treated with FGF-1 were compared by flow cytometry. Dot plots and histograms depict the results obtained in one representative donor; the grey histogram represent mock treated M2 macrophage, and the blue histogram represent the fluorescent profile of FGF-1 treated M2 macrophage stained with the indicated antibodies; the percentage and MFI (mean fluorescence intensity) of positive cells is indicated. The graph (lower panel) represents mean results from at least four different donors.

Following TTHX1114 treatment, we observed a significant reduction in the percentage of *in vitro* generated human monocyte-derived pro-inflammatory M1 macrophages expressing CD80, as detected by flow cytometry (**Figs. 7A**, *lower two panels*). Similarly, there was a significant reduction in the level of CD80 expressed on pro-inflammatory M1 macrophages following TTHX1114 treatment as measured by mean fluorescence intensity (MFI) (**Figs. 7A**, *lower two panels*). In contrast to pro-inflammatory M1 macrophages, TTHX1114 treatment led to a significant increase in the number of *in vitro*-generated human monocyte-derived anti-inflammatory M2 macrophages expressing CD206, as detected by flow cytometry (**Figs. 7B**, *lower two panels*).

Moreover, we investigated the signature of pro- and anti-inflammatory M1 and M2 cytokines secreted by M0/M1/M2 human monocyte-derived macrophages following TTHX1114 treatment using the Luminex detection platform (**Supplemental Fig. 4**). As shown in **Fig. 8**, TTHX1114 treatment had an overall trend in reduced production of pro-inflammatory cytokines (i.e., IL1α, IL-2, IL-12, IL-15, IL17-α, TNF-α, CCL-5) and chemokines (i.e., CXCL10) produced by the monocytes-derived pro-inflammatory M1 macrophages.

**Figure 8.**
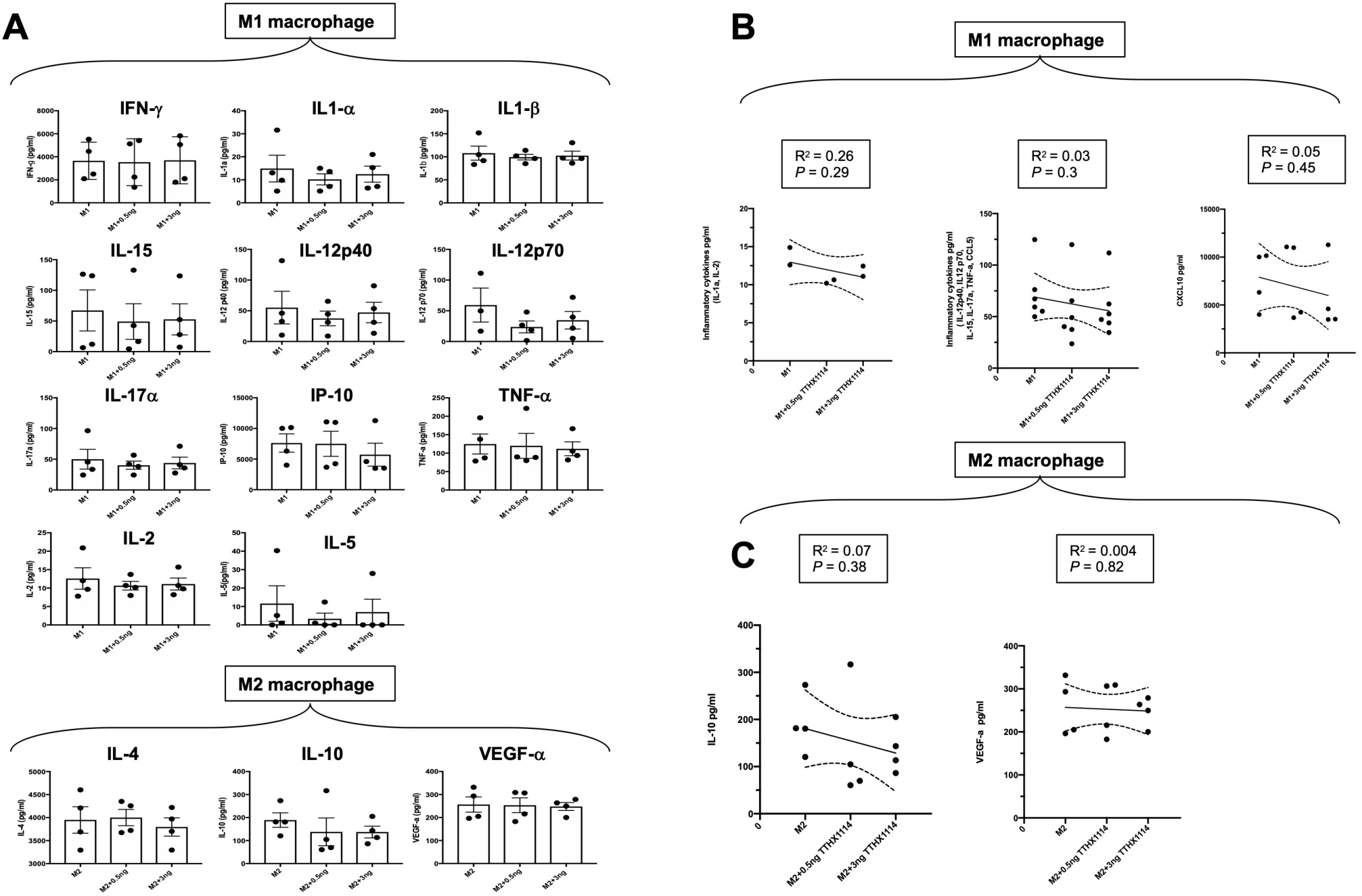
Effect of FGF-1 on cytokine secretion by human monocyte-derived M1 and M2 macrophages: Unpolarized M0 macrophages were generated from M-CSF-stimulated primary monocytes and then treated with FGF-1 (0.5 and 3 ng/ml) for 24 hr. M1- and M2-polarized macrophages were then generated by stimulation with IFN-γ and IL-4, respectively. (**A**) M1 signature cytokine (IFN-γ, IL-1α, IL-1β, IL-2, IL-5, IL-12p40, IL-12p70, IL-15, IL-17α, IP-10, TNF-α,) levels in the culture supernatants of M1 macrophage treated with FGF-1 and M2 signature cytokine (IL-4, Il-10, VEGF-α) levels in the culture supernatants of M2 macrophage treated with FGF-1. (**B**) Correlation graph showing level of M1 cytokines with FGF-1 dose kinetics. (IL-1α, IL-1β, IL-12p40, IL-12p70, IL-15, Il-17α, TNF-α, CCL5). (**C)** Correlation graph showing level of M2 cytokines with FGF-1 dose kinetics (IL-10, VEGF-α).

These findings in humans confirm that FGF-1 treatment skews polarization of monocyte-derived macrophages into the anti-inflammatory M2 phenotype. Moreover, FGF-1 treatment appeared to reduce production of anti-inflammatory mediators.

## DISCUSSION

Currently, one of the major unmet clinical challenges remains finding an effective medical treatment to offset vision-threatening inflammatory corneal herpetic disease. Corneal herpes simplex infections are among the most frequent serious ocular infections and a major cause of viral-induced blindness. Ocular infection with HSV-1 can cause eye disease ranging in severity from blepharitis, conjunctivitis, and dendritic keratitis, to disciform stromal edema and necrotizing stromal keratitis. To our knowledge, this is the first study to demonstrate that topical cornea treatment with FGF-1 has an anti-inflammatory role that reduces corneal keratopathy in a mouse model of primary and recurrent ocular herpes. The decreased frequency and function of pro-inflammatory macrophages M1 infiltrating the cornea was associated with reduced HSV-1-induced corneal immunopathology observed in both primary and recurrent ocular herpes.

Fibroblast growth factor-1 (FGF-1), a naturally occurring protein, promotes tissue repair and regenerates corneal tissue (23). The FGF family consists of a group of homologous growth-promoting polypeptides that increase proliferation, angiogenesis, and wound healing (24–26). Several studies have shown modulation of inflammatory responses by FGFs (25, 27–29). However, the role of FGF-1 on inflammation induced by herpes infection is not currently known. Trefoil’s engineered FGF-1 TTHX1114 builds on the well-known activities of naturally-occurring (native) FGF-1 to enable its use as a pharmaceutical for corneal diseases. Native FGF-1 is a potent stimulator of cell proliferation and migration, and has cell protective properties, all key attributes for its use in corneal disease treatment. The compound uniquely activates all seven forms of the FGF receptor, contributing to its potency. Unlike the naturally-occurring FGF-1 with an extremely short half-life, TTHX1114 is much more stable making it more suitable for pharmaceutical use.

HSV-1 infections of the cornea range in severity from minor transient discomfort to the blinding inflammatory disease herpes stromal keratitis (30). Here, we report a novel observation of anti-inflammatory effect of an engineered FGF-1 (TTHX1114) that healed both primary and recurrent corneal herpetic immunopathology leading to transparency of cornea, which is essential for normal vision. This anti-inflammatory role of engineered FGF-1 was associated with a decrease in the frequency and function of pro-inflammatory M1 macrophages infiltrating the cornea and, in contrast, an increase in the frequency and function ofanti-inflammatory M2 macrophages infiltrating the cornea. These results agree with a previous report by Dr. Rouse that similarly demonstrated the inhibition of VEGF signaling with a Src Kinase inhibitor ameliorated stromal keratitis (31) Although the anti-inflammatory role of FGF-1 is known in other disease conditions like renal diseases, its role in viral infection is not currently known. Nevertheless, it remains to be determined: (*i*) whether the FGF-1 treatment can accelerate healing of primary and recurrent corneal herpetic disease in humans; and (*ii*) a potential role of other innate and adaptive immune cells in the observed anti-inflammatory role FGF-1 in HSV-1-induced immunopathology.

Our present study is the first to demonstrate a reduction of corneal herpetic keratopathy following FGF-1 treatment associated with a decrease of cornea-resident pro-inflammatory macrophages M1. FGF-1 appeared to shift corneal-resident macrophages toward M2 phenotype. It remains to be determined whether HSV-1 replication in M1 and M2 macrophages was lowered following FGF-1 treatment. Moreover, we showed that the M1 macrophages expressed significantly lower levels of HSV-1-induced pro-inflammatory cytokines and chemokines following FGF-1 treatment. Thus, these findings shed significant light on a novel therapeutic approach to reducing primary and recurrent corneal herpetic disease by modulating both the phenotype and function of cornea-resident macrophages, which play a predominant role in the corneas following ocular herpes infection. We therefore suggest that inclusion of FGF-1 as a novel immunotherapeutic regimen against ocular herpes to skew cornea-resident macrophage development toward an anti-inflammatory M2 phenotype, rather than a pro-inflammatory M1 phenotype.

In the present report, the observed anti-inflammatory role FGF-1 in HSV-1-induced immunopathology was not associated with a significant effect on the frequency and activation of total CD4^+^ and CD8^+^ T cells that infiltrate HSV-1-infected corneas. The engineered FGF-1 (TTHX1114) affects the function and frequency HSV-specific CD8^+^ T cells, but not HSV-specific CD4^+^ T cells, that infiltrate HSV-1-infected corneas with a yet-to-be determined mechanism. Thus, the effect of FGF-1 on cornea-resident HSV-specific CD8^+^ T cells remains to be determined, since CD4^+^ T cells appear to be the main orchestrators of the blinding immunoinflammatory lesion that represents an immunopathological response to HSV-1 infection. Moreover, corneal herpetic lesions have an increased severity if the regulatory Foxp3^(+)^CD4^+^ Treg response is compromised from the onset of infection (32). Tregs beneficially influence the severity of ongoing tissue-damaging immune responses to HSV-1 infection (32). This suggest that therapies, such as FGF-1, boosting Treg function in the clinical phase hold promise for controlling a lesion that is an important cause of human blindness. A potential effect of FGF-1 on cornea-resident Foxp3^(+)^CD4^+^ Treg must be determined during the ongoing anti-inflammatory effect of FGF-1.

The underlying cellular and molecular mechanisms that led to FGF-1 treatment decreasing HSV-1-induced corneal immunopathology remain to be determined. HSV-1 infection of the cornea induces lymphangiogenesis that continues to develop well beyond the resolution of infection. In this report, we discovered that topical cornea treatment with FGF-1 has an anti-inflammatory role that reduced corneal keratopathy in a mouse model of primary and recurrent ocular herpes. The decrease in the frequency and function of cornea-resident pro-inflammatory macrophages M1 was associated with reduced HSV-1-induced corneal immunopathology observed in both primary and recurrent ocular herpes following FGF-1 treatment.

Excessive proteolysis and neovascularization, resulting in corneal scarring has been associated with loss of corneal clarity. Inflammatory leukocytes infiltrate the cornea and have been implicated to be essential for corneal neovascularization, a clinically relevant manifestation of stromal keratitis. Cornea infiltrating leukocytes including neutrophils and T cells appear to not have a significant role in corneal neovascularization past virus clearance. Multiple pro-angiogenic factors, including FGF-1, are expressed within the cornea following virus clearance. Many angiogenic factors such as vascular endothelial growth factors are present in the cornea but their angiogenic activities are impeded by being bound to a soluble form of the VEGF receptors. It is likely that an imbalance between vascular endothelial growth factors and their receptors present in the cornea occur after ocular HSV-1 infection may cause prominent neovascularization, an essential step in the pathogenesis of the vision-impairing lesion, stromal keratitis. However, the significant effect of FGF-1 on HSV-1-induced corneal immunopathology in the B6 mouse model of primary ocular herpes was independent of lymphangiogenesis. Indeed, the immunohistochemistry and FACS analysis revealed that FGF-1 treatment did not affect lymphangiogenesis and lymphocyte infiltration in HSV-1-infected mice.

In conclusion, we report here that compared to HSV-1 infected non-treated mice, the infected and FGF-1 treated mice showed (i) an overall resistance to disease; (ii) a significant decrease in primary stromal keratitis (days 5, 14, and 21) and blepharitis (days 7 and 14); (iii) a significant decrease in disease duration in herpes reactivation and (iv) a significant decrease in corneal inflammatory macrophage. The effect of FGF-1 seen on mouse macrophages was conformed on human macrophage pointing to a potential clinical application. However, FGF-1 treatment did not affect the number and function of cornea resident T cells nor virus corneal replication. Topical cornea treatment with eFGF-1 is associated with reduced corneal keratopathy in a mouse model of primary ocular herpes. Increased frequency and function of anti-inflammatory M2 macrophages was associated with reduced corneal keratopathy observed in the FGF-1 treated cornea.

## ACKNOWLEDGEMENTS

This work is supported by a grant from Trefoil Therapeutics, Inc. and by Public Health Service research grants EY019896, EY14900 and EY024618 from the National Eye Institutes (NEI) and AI150091, AI143348, AI147499, AI143326, AI138764, AI124911 and AI110902 from the National Institutes of Allergy and Infectious Diseases (NIAID) to LBM and from The Discovery Center for Eye Research, and Research to Prevent Blindness.

